# Last and corresponding authorship practices in ecology

**DOI:** 10.1101/126938

**Authors:** Meghan A. Duffy

## Abstract

Authorship is intended to convey information regarding credit and responsibility for manuscripts. However, while there is general agreement within ecology that the first author is the person who contributed the most to a particular project, there is less agreement regarding whether being last author is a position of significance and regarding what is indicated by someone being the corresponding author on a manuscript. Using an analysis of papers published in *American Naturalist*, *Ecology*, *Evolution,* and *Oikos,* I found that: 1) the number of authors on papers is increasing over time; 2) the proportion of first authors as corresponding author has increased over time, as has the proportion of last authors as corresponding author; 3) 84% of papers published in 2016 had the first author as corresponding author; and 4) geographic regions differed in the likelihood of having the first (or last) author as corresponding author. I also carried out an online survey to better understand views on last and corresponding authorship. This survey revealed that most ecologists view the last author as the “senior” author on a paper (that is, the person who runs the research group in which most of the work was carried out), and most ecologists view the corresponding author as the person taking full responsibility for a paper. However, there was substantial variation in views on authorship, especially corresponding authorship. Given these results, I suggest that discussions of authorship have as their starting point that the first author will be corresponding author and the senior author will be last author. I also suggest ways of deciding author order in cases where two senior authors contributed equally.

## Introduction

Who is the last author on a paper? Depending on authorship conventions in a field, the last author might be the person whose surname comes last alphabetically, the person who runs the research group where the research was done, or simply the person who did the least work on the project (Tscharntke et al. 2007). In math, for example, authorship tends to be determined alphabetically (Waltman 2012), whereas in biomedical fields, the last author position is one that tends to carry extra weight (Moulopoulos et al. 1983, Wren et al. 2007, Venkatraman 2010). In ecology, alphabetical author lists are not the norm, but standard authorship practices have received relatively little study. Thus, we are in a similar situation to the one described in 1997 by Rennie et al. when they discussed order of authorship and what it conveys: “Everyone is equally sure about their own system; the point is that none of these schemes is actually disclosed, so the readers, to whom this should be addressed, are not let in on the secret: they have not been told which code book to use and how it works.” The goals of this study are to see if the number of authors and the position of the corresponding author have changed over time, to describe the current systems in use by ecologists regarding last and corresponding authorship, and to see whether certain factors (e.g., research area, career stage) are associated with views on authorship.

As noted in an earlier publication on this topic (Tscharntke et al. 2007), the first author of an ecology paper is generally the person who made the greatest overall contribution to the work, but there is no consensus on how to determine the order of the remaining authors. In a survey of 57 ecologists at the 2004 meeting of the Ecological Society of America, respondents gave ten unique authorship order combinations for a scenario involving only three potential coauthors, with respondents disagreeing about both who should be included as an author and the order of authorship (Weltzin et al. 2006). There is also confusion over what is signified by corresponding authorship (Laurance 2006).

This is problematic for two reasons. First, people are assessed based on their publication records, meaning that unclear authorship criteria make it difficult to determine how much credit an author should get for a publication (Tscharntke et al. 2007, Wren et al. 2007, Eggert 2011). Job applications, grant proposals, and tenure and promotion decisions are all impacted by publication records. If people evaluating these applications, proposals, and dossiers have different views on what it means to be last or corresponding author, then authorship order does not provide a reliable signal. This can be problematic if, for example, an assistant professor puts herself as last author as an indicator of having led the work, but a tenure letter writer thinks she is last because she did the least work. Second, authorship on a publication entails not just credit for the work, but responsibility for it as well (Rennie et al. 2000, Venkatraman 2010, Eggert 2011). In cases where concerns about research are raised, it is important to know, for example, if corresponding authorship indicates that someone is taking full responsibility for the publication.

In this study, I first present data on the number of authors over time as well as the position of the corresponding author over time in four journals (*American Naturalist, Ecology*, *Evolution,* and *Oikos*). For papers published in these four journals in 2016, I also asked whether geographic region or number of authors influenced the likelihood of having the first (or last) author as corresponding author. I also present results of a survey of scientists (80% of whom identified ecology as their primary research area) that asked about views on last and corresponding authorship. In addition to giving information on overall views of ecologists, the survey allowed me to explore whether factors such as research subfield, time since PhD, geographic location, and amount of interdisciplinary work were associated with views on last and corresponding authorship. I end by suggesting that, since most readers expect authors to use a first-last author emphasis (FLAE, sensu Tscharntke et al. 2007) and since the vast majority of papers in *American Naturalist, Ecology*, *Evolution,* and *Oikos* have the first author as the corresponding author, those are good starting places for discussions regarding author order and corresponding authorship (while recognizing that there will be situations where it is desirable or necessary to deviate from this). I also give suggestions for how to determine authorship order in cases where two “senior” authors have made equal contributions to a study.

## Methods

### Literature survey

The literature survey involved a combination of approaches. First, I began by reviewing the first issue of the journal *Ecology* every ten years from 1956-1986. In 1996, I reviewed the second issue of the journal, since the first contained a special feature and I wished to avoid any potential confounding effects of analyzing a special feature. I used this data set to look at corresponding authorship practices in *Ecology* from 1956-1996, tracking whether there was a note indicating to whom correspondence (or reprint requests) should be sent. Second, I collected data from Web of Science on the number of authors of papers published in all issues of *Ecology* every ten years from 1956-1996 and every five years from 2001-2016, as well as from the journals *American Naturalist*, *Evolution*, and *Oikos* every five years from 2001-2016. Third, for 2001, 2006, 2011, and 2016, I also extracted data on corresponding authorship from Web of Science. I considered authors who had their email addresses in the Web of Science record as corresponding authors (but see note below about exceptions, especially in 2001&2006). Corresponding authorship was then grouped into six categories: 1) “first” (the email address given was for the first or only author in the author string), 2) “middle” (the email address given was for someone other than the first or last author), 3) “last” (the email address given was for the last author), 4) “ND” (not designated; when an email address was not given for any author), 5) “all” (when both – for papers with only two authors – or all of the authors on a paper had email addresses given), and 6) “other” (when email addresses were given for some other combination of authors, such as the first and last). For one paper in *Oikos*, an email address was given but it was not possible to determine which author the email address corresponded to; this paper was omitted from the analysis. For all four journals in 2001 and for *American Naturalist* in 2006, the email addresses given (or not given) by Web of Science did not match what appeared on the first page of the article in the print journal. In most cases in 2001, the issue was the omission of email addresses; for *American Naturalist* in 2006, the issue was that Web of Science had email addresses for all authors in most cases, whereas the print copy indicated one author for correspondence. Thus, for all four journals in 2001 and for *American Naturalist* in 2006, I did not use Web of Science data regarding corresponding authorship. Instead, I manually reviewed the papers in the first 900 pages of each journal in that year to determine corresponding authorship, using the same criteria given above. (This was done by visiting the stacks in the University of Michigan library; Figure 1.) In some cases, email addresses were given for multiple authors but one author was indicated as the one to whom correspondence should be addressed; in these cases, only the author designated for correspondence was considered the corresponding author. Editorial material, book reviews, retractions, and corrections were excluded from analyses.

**Figure 1.**
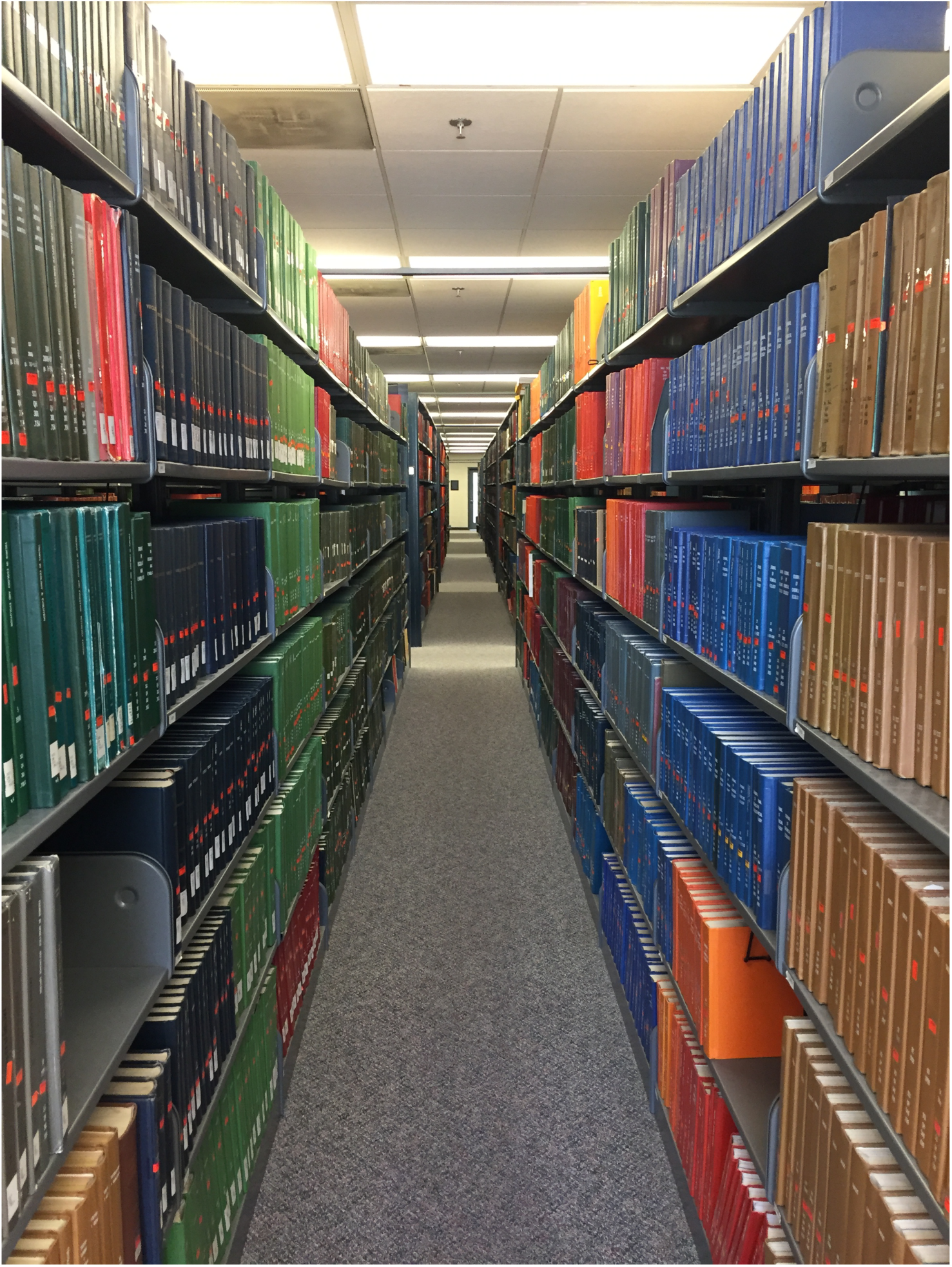
Stacks containing bound volumes of journals (Shapiro Library, University of Michigan)

For the journal *Ecology*, changes in the number of authors over time (1956-2016) were analyzed using a glm with Poisson error. For 2001-2016, I used the dataset on number of authors from all four journals and a glm (again, with Poisson error) with year, journal, and their interaction as fixed effects. Changes (over 2001-2016) in whether the first author was corresponding author were analyzed using a glm with binomial error with year, journal, and their interaction as fixed effects. This analysis was also carried out for whether the last author was corresponding author.

For the 2016 publications, I also extracted information on where the reprint author lived, and used that to compare corresponding authorship by region using a glm with binomial error and logit link function. In most cases, there was only one reprint author indicated; however, in cases where there were multiple addresses, I used the country indicated in the last address. The regions used in this analysis were Africa, Asia, Europe, North America (which included Canada, Jamaica, Mexico, Panama, and the United States), Oceania (which included Australia and New Zealand), and South America. I did this analysis once with a response variable indicating whether the first author was the corresponding author, and once with a response variable indicating whether the last author was the corresponding author. For the statistical analysis, I only included regions with at least 50 publications (that is, Asia, Europe, North America, and Oceania).

I also looked at whether the number of authors influenced whether the first or last author was the corresponding author; because of the small sample sizes for papers with 10 or more authors, I combined papers with 10 or more authors and treated the number of authors as an ordinal predictor. This analysis used data from all geographic regions, but omitted papers with only one author (as those could not have a last author as corresponding author, based on the authorship definitions I used).

All analyses were carried out in R (v 3.4.1). Figures were also made in R using the ggplot (Wickham 2009) and cowplot (Wilke 2017) packages. Data and code for the analyses and figures are available at: http://github.com/duffymeg/DEAuthorshipPoll

### Poll

I carried out a poll of readers of the *Dynamic Ecology* blog. In addition to appearing on the blog, the poll was advertised via social media and thus likely reached a wider readership than a typical blog post. The poll first appeared on 6 April 2016 and ran for two weeks. After removing four blank responses, there were 1122 responses to the poll.

The poll had four main questions: 1) For ecology papers, do you consider the last author to be the senior author? 2) Which of the following statements most closely matches the current norms in ecology in terms of who is corresponding author? 3) Which of the following statements would be best practice in terms of who is corresponding author? and 4) If someone includes a statement on his/her CV indicating they have used a first/last author emphasis, do you pay attention to that? The poll also asked about the respondent’s primary research area, whether their research is primarily basic or applied, how frequently they conduct interdisciplinary research, how many years post-PhD they are, where they live (options: Africa, Asia, Australia, Europe, North America, and South America), and what their current department is (divided by discipline: EEB, biology, natural resources, or other). The full survey, including the questions and all the answer options, is given in the Supplement.

In addition to presenting the overall responses to the four main questions, I used the additional information on research area, geographic location, years since degree, and department type to look for factors associated with views on last and corresponding authorship. Prior to doing those analyses, I decided that a difference between two groups in their views on authorship had to be at least 10% in order to be considered notable. While this threshold is somewhat arbitrary, it helped ensure that small differences weren’t overinterpreted. Data were analyzed in R (v 3.4.1) and plotted using the ggplot (Wickham 2009), cowplot (Wilke 2017), and likert (Bryer and Speerschneider 2016) packages. For the analysis of views on last authorship, responses were turned into a binary response based on whether they viewed the last author as likely to be the senior author (with “Yes”, “It depends, but probably yes”, and “Not sure, but probably yes” all being coded as 1 and the other three responses as 0). For the analysis of views on current corresponding authorship practices, I created a binary variable based on whether someone chose the “full responsibility” option (that is, whether or not they chose the option saying that the corresponding author “uploaded the files, managed the revisions and wrote the response to reviewers, and took responsibility for the paper after publication”).

For the analysis of differences across career stages, I excluded data from the 19 respondents who did not have PhDs and were not in graduate school, then treated the other categories as ordinal variables and looked a linear effect of career stage (years since PhD) on views on last or corresponding authorship. For analyses related to geography, I compared views of people currently living in Europe with those of people currently living in North America. For analyses related to research area, I compared responses of people who identified primarily as ecologists with those of people who identified primarily as evolutionary biologists. For analyses of department type, I compared responses of people who are in EEB departments with those of respondents in Biology and Natural Resources departments. Finally, for the analysis of views on last authorship, I also tested for effects of whether someone primarily does basic or applied research, and of the frequency with which they do interdisciplinary research (modeled as an ordinal variable). Neither basic vs. applied research nor the amount of interdisciplinary research significantly influenced views on last authorship; therefore, in the interest of space, those results are not presented below. All analyses were done using glms in R with binomial error and a logit link function.

One important caveat for this study, as discussed further in the discussion section, is that there are surely biases related to this being a voluntary, online poll of blog readers. Among other things, the poll respondents are likely to be younger, on average, than ecologists as a whole. One conclusion of this study is that this area would benefit greatly from additional study by social scientists with formal training in survey design and qualitative analysis.

Aside from expecting the number of authors on papers to increase over time (as has been found by others: Johnson 2006, Weltzin et al. 2006, Fox et al. 2016, Logan 2016), I did not have strong *a priori* hypotheses about how corresponding authorship patterns would change, or about whether or how research area, geographic location, years since degree, and department type might influence patterns of corresponding authorship or views on last and corresponding authorship.

## Results

### Authorship over tim

The number of authors on *Ecology* papers is increasing over time (*Z* = 24.46, *p* < 0.0001), with a particularly notable uptick after 1996 (Figure 2A). In 1956, the median number of authors on a paper was 1 (mean = 1.4), whereas in 2016 the median was 4 (mean = 4.6). Between 1956 and 1996, the corresponding author on a paper was not usually indicated and mailing addresses for all authors were given. Of the 129 papers analyzed during that window, only two indicated the author to whom correspondence should be addressed; in other words: it was very rare for a corresponding author to be indicated during this time window. Interestingly, in one of the cases (Kalisz and Teeri 1986) the first author was indicated, whereas in the other (Murcia and Feinsinger 1996) the second author was indicated.

**Figure 2.**
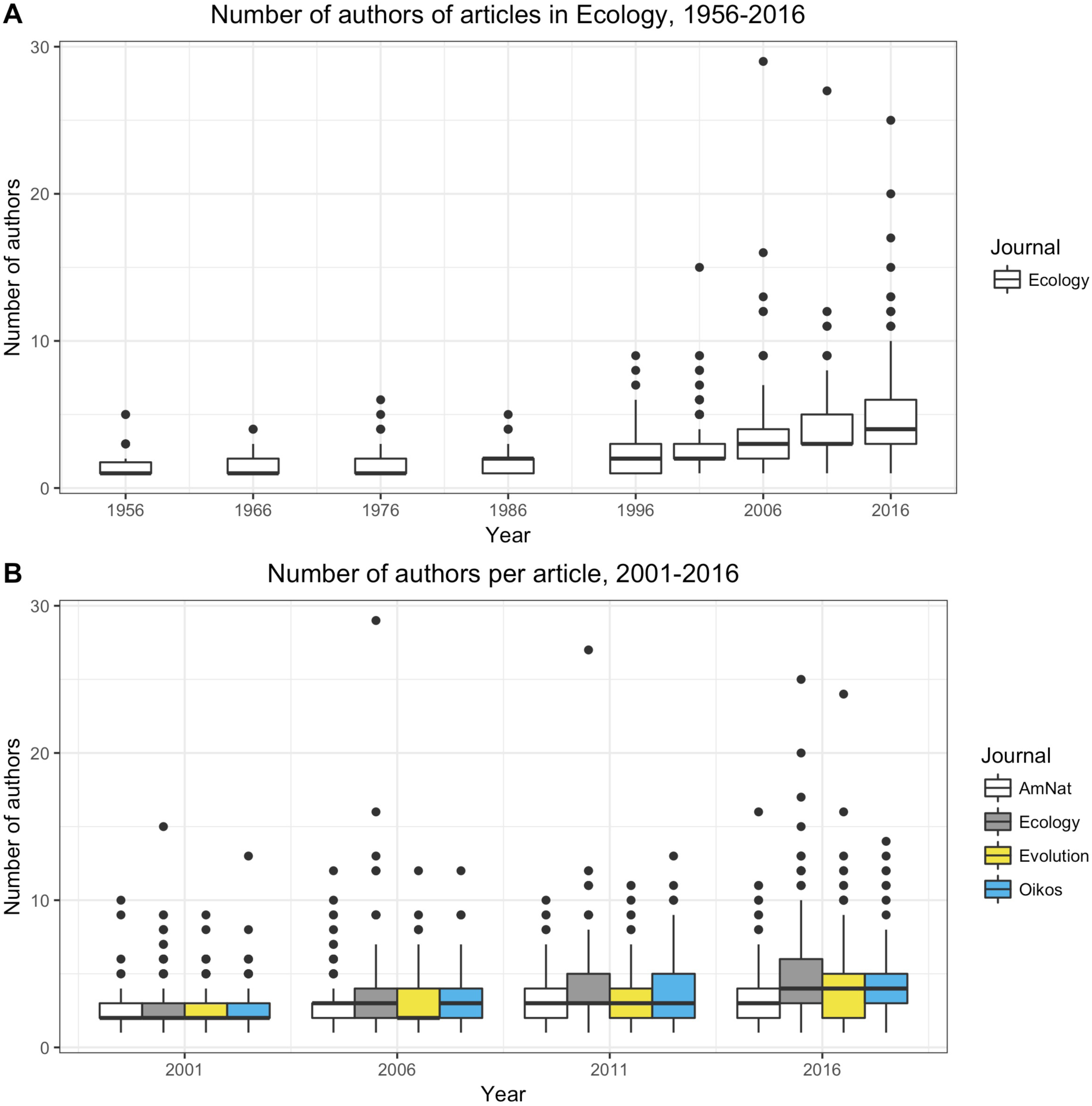
Number of authors on papers in *American Naturalist, Ecology*, *Evolution,* and *Oikos* over time. See methods for more information on which journal issues were analyzed. A) Data for *Ecology* for 1956-2016. B) Data for *American Naturalist*, *Ecology*, *Evolution,* and *Oikos* for 2001-2016.

Looking across all four journals for the period 2001-2016, the number of authors increased over time (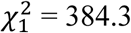, p < 0.0001; Figure 2B) and journals differed in the number of authors per paper (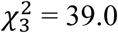, p < 0.0001), but there was not a significant difference between journals in the increase in the number of authors over time (that is, there was not a significant journal*year interaction: 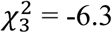, p = 0.097).

The proportion of first authors as corresponding author increased over time (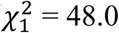, p < 0.0001; Figure 3) and differed between journals (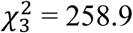, p < 0.0001); moreover, the change in first author as corresponding author over time differed between journals (interaction: 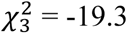, p = 0.0002). *American Naturalist* and *Evolution* showed high proportions of papers with all authors having email addresses in 2001 and 2006, whereas this was rare in all journals in 2016 (Figure 3). The proportion of last authors as corresponding author also increased over 2001-2016 (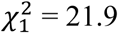, p < 0.0001; Figure 3); the proportion of last authors as corresponding author did not differ significantly between journals (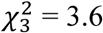, p = 0.31), nor did journals differ significantly in the increase over time (interaction: 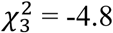, p = 0.19). In 2016, the corresponding author was usually the first author (range across the four journals: 77-90% of papers); less commonly, it was the last author (range across the four journals: 9-18% of papers).

**Figure 3.**
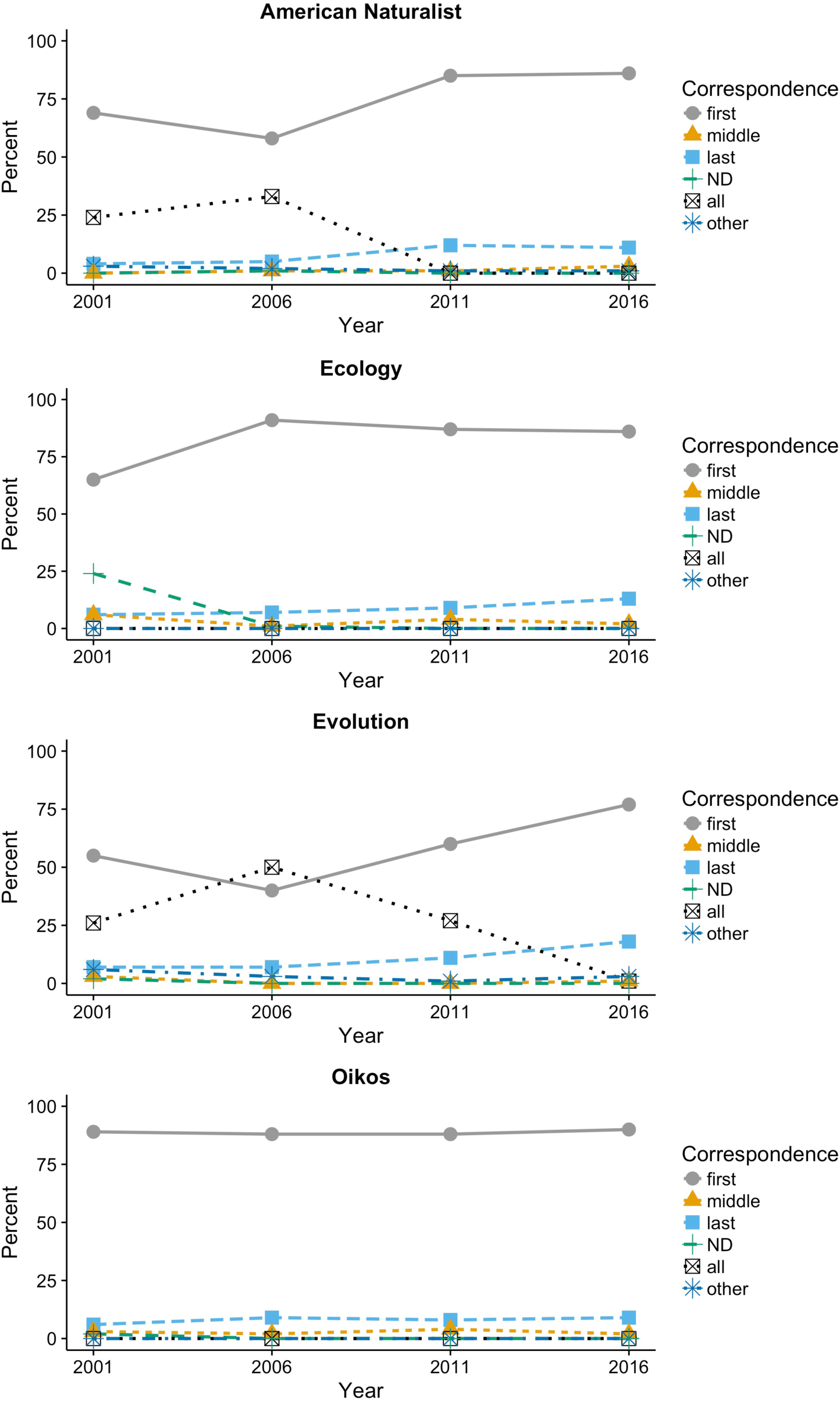
Corresponding author position for articles in *American Naturalist*, *Ecology*, *Evolution,* and *Oikos*. “ND” means that a corresponding author was not designated.

### Analysis of corresponding authorship in 201

Geographic regions differed in the likelihood of having the first (or last) author as corresponding author. Focusing on the regions with at least 50 publications in the dataset, papers where the reprint author lived in Asia were much less likely to have the first author as corresponding author (Figure 4A; pairwise comparisons to Europe, North America, and Oceania: all *Z* > 3.1, all *p* < 0.002) and more likely to have the last author as corresponding author (all *Z* < −2.7, all *p* < 0.006). Papers where the reprint author lived in Europe were less likely to have the first author as corresponding author than ones where the reprint author lived in North America (*Z* = −1.99, *p* = 0.047), but this effect was more modest (83% vs. 88%).

**Figure 4.**
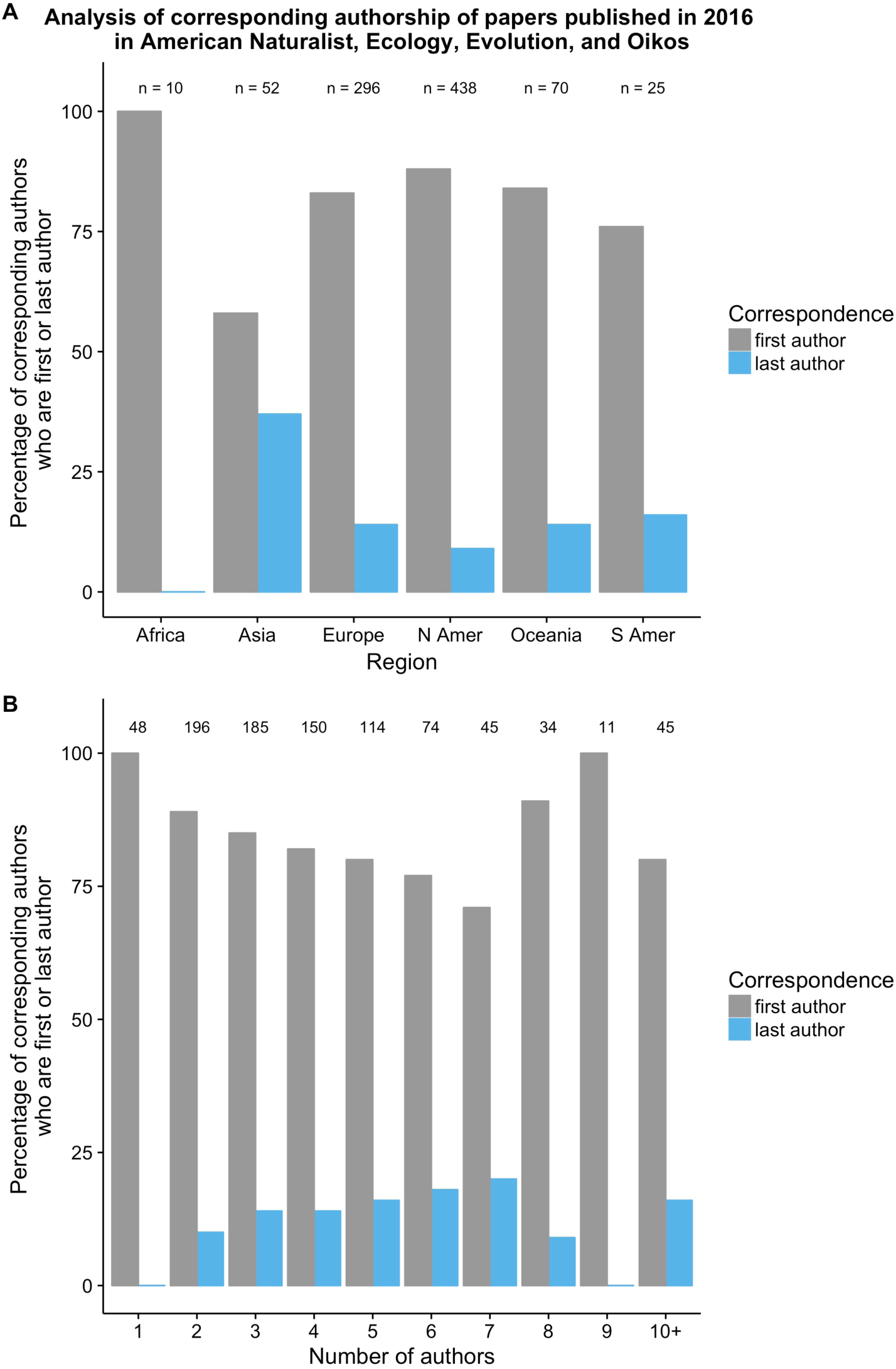
Influence of geographic region and number of authors on corresponding authorship. A) Percentage of corresponding authors from different geographic regions who are first author (gray bars) or last author (blue bars). The statistical analysis of this dataset only included regions with at least 50 publications. B) Relationship between the number of authors on a paper and whether the corresponding author is the first author (gray bars) or last author (blue bars). Numbers over the bars indicate the number of papers in that category. The gray and blue bars do not always sum to 100% because, rarely, the corresponding author was a middle author or a combination of authors (see Figure 3 for general patterns).

There was no clear relationship between the number of authors on a paper and the likelihood of the corresponding author being first (linear regression term for model with 10 or more authors binned together: *Z* = 0.032, *p* = 0.975) or last (linear regression term: *Z* = −0.031, *p* = 0.975) author (Figure 4B). If papers with 7 or more authors were binned together, there was still not a significant effect of number of authors on last authorship (linear regression term: *Z* = 1.59, *p* = 0.11), but there was a significant effect on first authorship (linear regression term: *Z* = − 2.53, *p* = 0.012).

### Demographics of poll respondent

80% of respondents indicated that ecology was their primary research field (Table 1). Most poll respondents were current students (28%) or had received their PhD within the past 1-5 years (31%), but respondents included people in all categories, including those who received their PhD over 20 years ago (Table 2). The vast majority of the poll respondents live in North America (64%) or Europe (26%; Table 3).

**Table 1.**
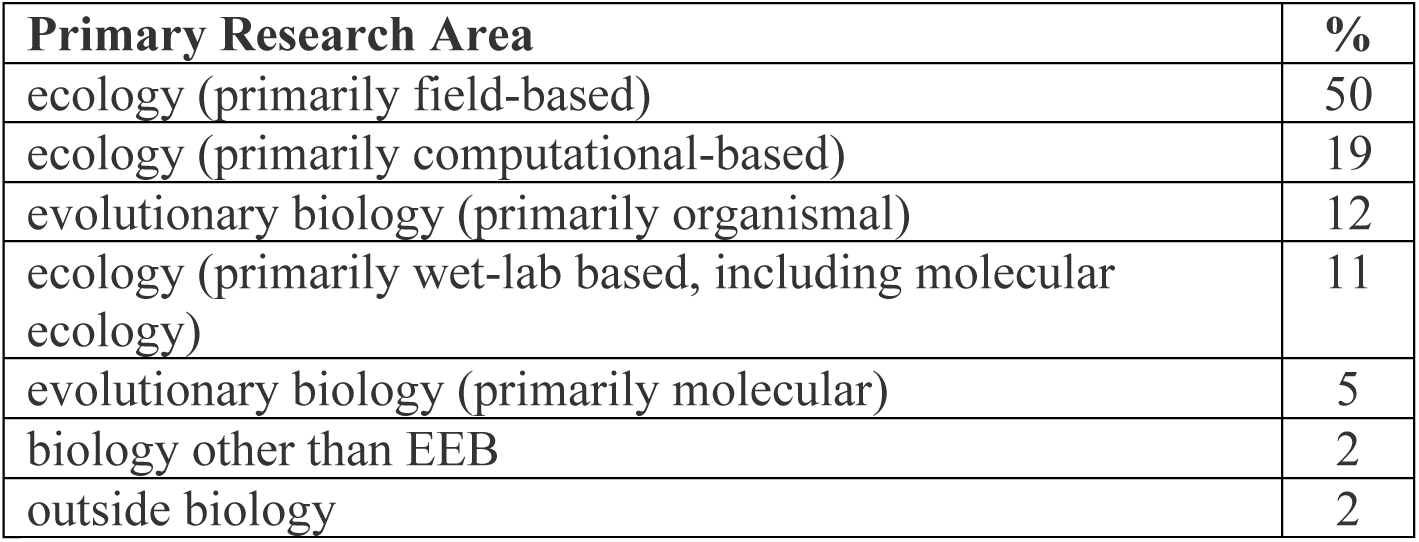
Primary research area of respondents to poll on last and corresponding authorship, sorted in decreasing order of commonness.

**Table 2.** Number of years since receiving PhD for poll respondents.

| Years since PhD | % |
| --- | --- |
| 0 (current students should choose this) | 28 |
| 1-5 | 31 |
| 6-10 | 18 |
| 11-15 | 12 |
| 16-20 | 5 |
| >20 | 5 |
| no PhD and not a current student | 2 |

**Table 3.**
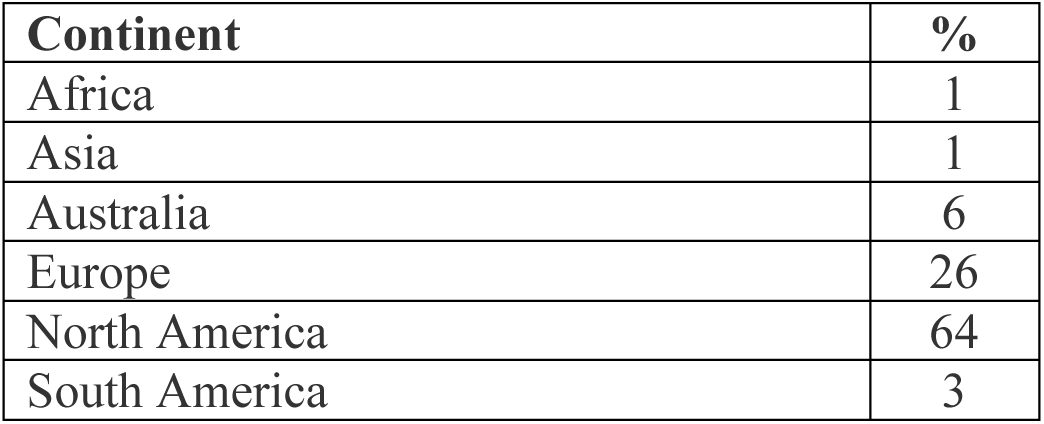
Geographic location of poll respondents, sorted alphabetically.

### Views on last authorship

For ecology papers, most respondents viewed the last author as the senior author (that is, the lab head or principal investigator; Figure 5A). However, this view is not unanimous: the three “no”-related answers garnered 14% of the responses. Confusion about whether the last author is the senior author could be reduced if ecologists included a note on their CV indicating that the last author position is one of emphasis. However, the poll results suggest this is likely to only be partially effective – 29% of respondents said they do not or would not pay attention to these statements (Figure 5B).

**Figure 5.**
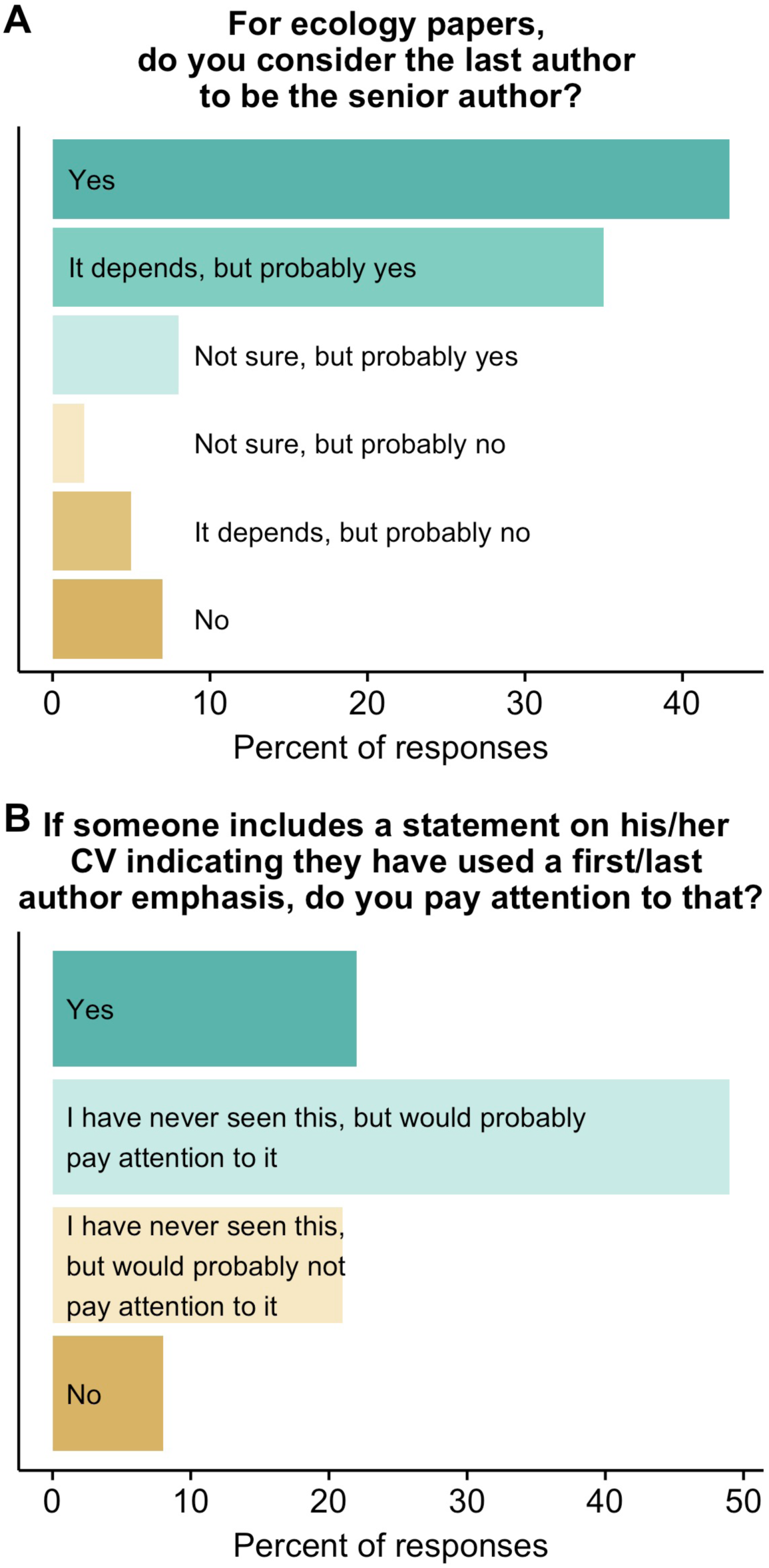
Views of poll respondents on A) whether the last author of a paper is the senior author and B) whether they would pay attention to a statement on the CV indicating that the last author position was one of emphasis.

Year of degree (as a proxy for career stage) influenced views on last authorship (Figure 6A), with people who are within 10 years of their PhD (or currently in graduate school) more likely to view the last author as senior author (as evidenced by a significant linear term in the regression: *Z* = −2.2, 0.028). Respondents living in Europe were more likely to say the last author is the senior author, as compared to those in North America (95% “yes” responses vs. 82%, respectively; *Z =* 5.3, *p* < 0.0001; Figure 6B). Looking at primary research area, the two evolution categories had the highest proportion of positive responses to the question about whether the last author was the senior author, with ecologists being somewhat less likely to give one of the “yes” responses (as compared to evolutionary biologists; Figure 6C; contrast of ecology vs. evolution: *Z* = 2.4, *p* = 0.02). People in Biology and EEB departments were more likely to view the last author as the senior author, compared to those in Natural Resources departments or other types of departments (Figure 6D; contrasts of EEB departments to Biology (*Z* = 0.23, *p* = 0.82), Natural Resources (*Z* = 3.03, *p* = 0.002), and other departments (*Z* = 3.22, *p* = 0.001)).

**Figure 6.**
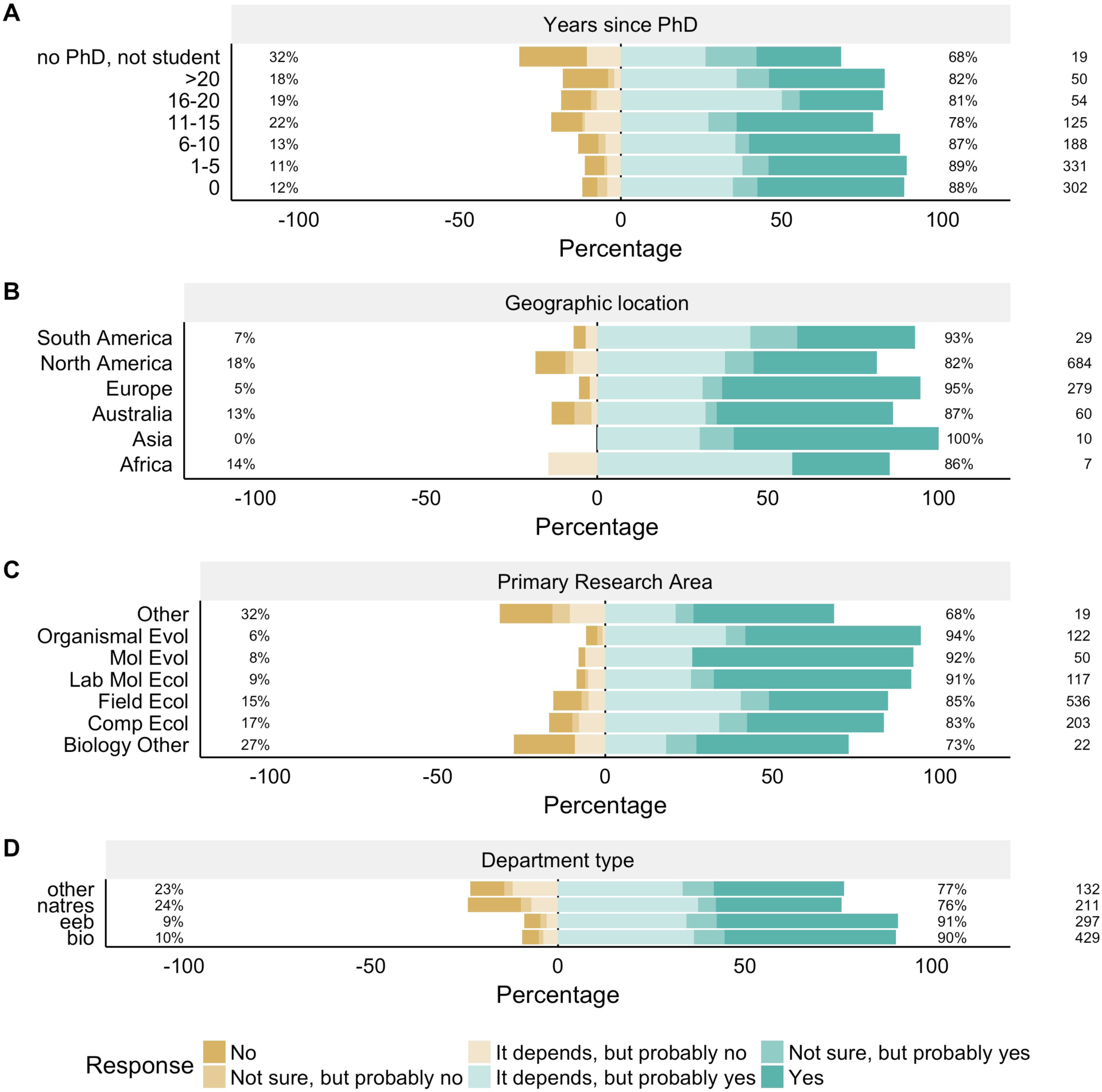
Variation in views on last authorship by career stage, geographic location, research area, and department type. The bars shaded in greens are positive responses to the question “For ecology papers, do you consider the last author to be the senior author”, whereas gold responses are negative responses (as described in the figure legend). The percentage on the right gives the total percentage of positive responses, while the percentage on the left gives the total percentage of negative responses for a group. The number on the right hand side shows the number of respondents in a given category (e.g., 29 respondents indicated that they live in South America).

### Views on corresponding authorship

There was substantial variation in respondents’ views on current and best practices for corresponding authorship (Figure 7). Most respondents (54%) said that the corresponding author “uploaded the files, managed the revisions and wrote the response to reviewers, and took responsibility for the paper after publication”. The next most common response (19% of respondents) was that the current practice is that the corresponding author is the person who simply uploaded the files – though only 8% viewed this as best practice. Only 7% said that the current practice is that the corresponding author is the senior author.

**Figure 7.**
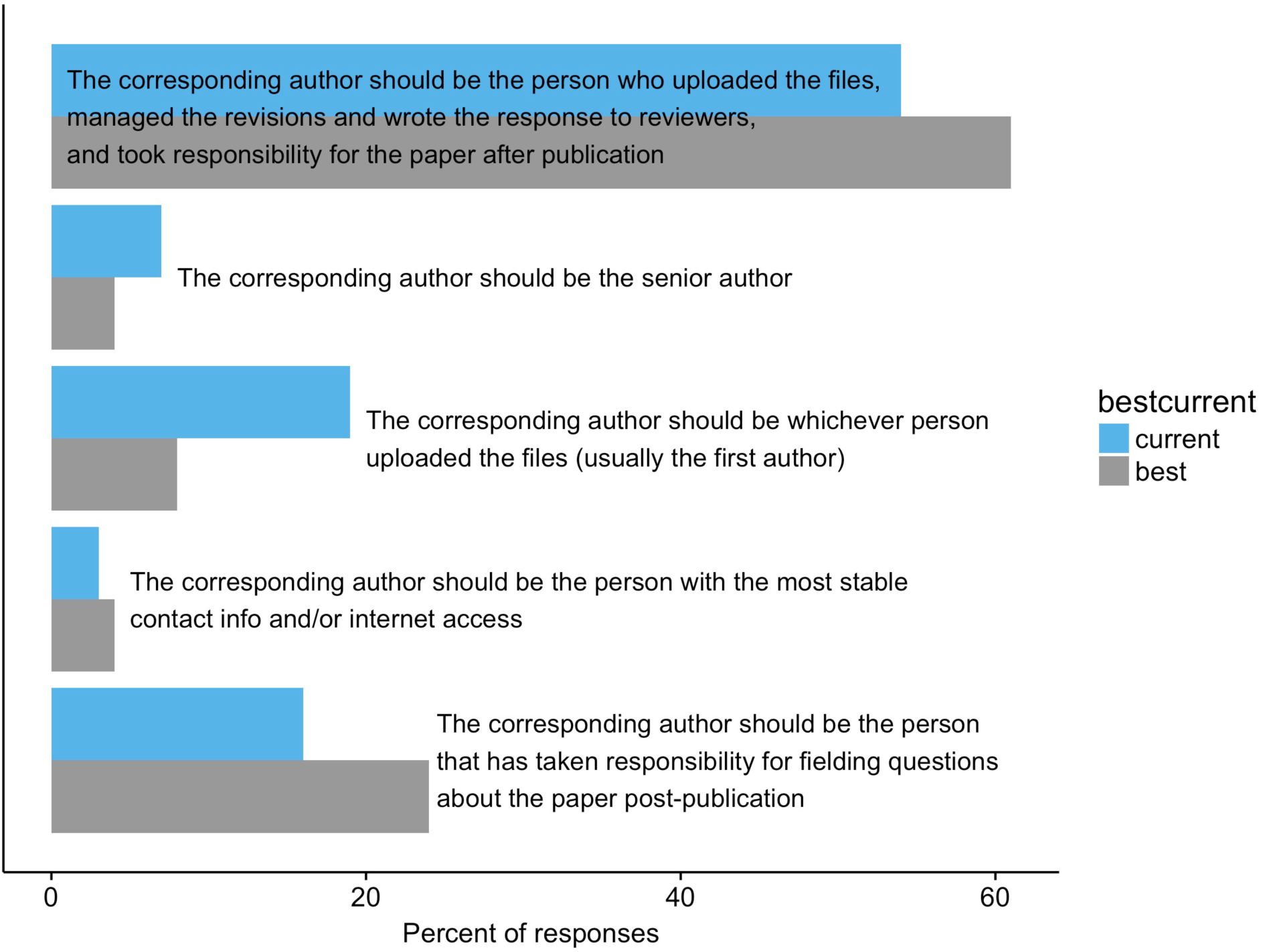
Views of poll respondents on current (light blue) and best (gray) practices for corresponding authorship.

Looking at the effects of career stage (that is, years since PhD), research area, department type, and geographic region on views on corresponding authorship practices, the only factor that was statistically significant and reached the 10% effect size threshold was department type (Figure 8): people in EEB departments were more likely to choose the “full responsibility” option (that is, to say the corresponding author “uploaded the files, managed the revisions and wrote the response to reviewers, and took responsibility for the paper after publication”) than those in Biology departments (60% vs. 50%, respectively; *Z* = 2.4, p = 0.016). There was no significant impact of career stage (linear regression term: *Z* = −1.3, *p* = 0.20), nor were there significant differences in ecologists vs. evolutionary biologists (*Z* = 1.12, p = 0.26) or those living in Europe vs. North America (*Z* = 1.6, p = 0.10).

**Figure 8.**
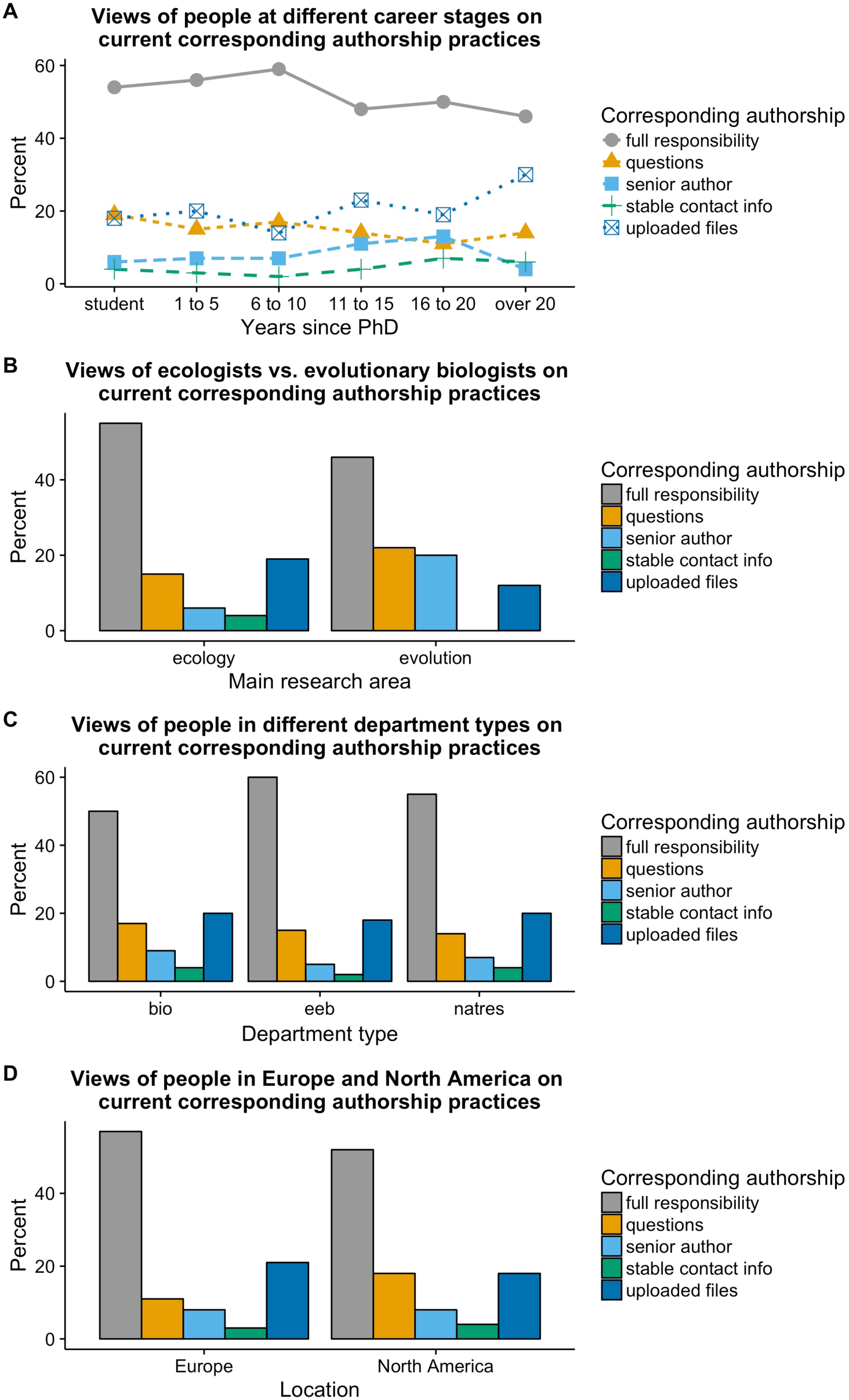
Influence of career stage, research area, department type, and geographic location on views on current corresponding authorship practices.

## Discussion

The number of authors on papers in ecology has increased over time; in 1956, most *Ecology* papers had only a single author, whereas in 2016 the median number of authors was 4. Prior to the late 1990s, it was rare for the corresponding author of a paper to be designated; now, the first author is usually the corresponding author, with the last author being the corresponding author in a minority of cases. Most ecologists view the last author as a position of emphasis in a paper, though this view is not universal. Most ecologists view the corresponding author as the person taking full responsibility for a paper, but, again, the survey revealed variation in views regarding current and best practices for corresponding authorship. Overall, there is variation in views on corresponding and last authorship in ecology, and the field would benefit from greater consensus on what is signified by corresponding and last authorship, as well as additional studies into the factors that influence decisions regarding corresponding and last authorship.

To state the obvious, decisions about who should be last and/or corresponding author are only necessary if there is more than one author. Thus, the trend in ecology towards having more authors on papers (Figure 2), as also seen by others (Johnson 2006, Weltzin et al. 2006, Fox et al. 2016, Logan 2016), means that there are more decisions to be made regarding authorship, including last and corresponding authorship.

Over the past several decades, various systems for attempting to indicate how much different authors contributed to multi-author papers have been proposed (e.g., Davis and Gregerman 1969, Moulopoulos et al. 1983, Rennie et al. 1997, Weltzin et al. 2006). A common suggestion is to use author contribution statements (e.g., Moulopoulos et al. 1983, Rennie et al. 1997, Cozzarelli 2004). While author contribution statements do have the potential to remove ambiguity about whether the last author is a position of emphasis, they have several problems themselves. First, unless the full author contribution statements are put on a CV for every publication, people reviewing job, grant, or award applications are unlikely to see them (especially at earlier stages of screening). Second, and more problematically, people do not necessarily trust author contribution statements (Venkatraman 2010, Fox 2016): in a different poll done on the Dynamic Ecology blog, only 41% of respondents indicated that author contribution statements are always or usually accurate in their experience (Fox 2016). One possible modification would be to make the author contribution statements less fine-grained: rather than indicating which authors carried out which specific tasks, contribution statements could indicate which research groups led different aspects of the project (e.g., “the X Lab led the empirical components of this work, and the Y Group led the development of the mathematical model”).

Thus, for the foreseeable future, people will continue to attempt to infer the contributions of different authors based on the order of authorship. The results of this survey demonstrate that, at present, most ecologists tend to view the last author as the senior author (Figure 5). Therefore, when discussing authorship, ecologists should assume that most people will interpret authorship order assuming a first-last author emphasis (FLAE), viewing the last author as the senior author. As a result, I recommend that discussions regarding authorship should have as their starting point that the senior author will be the last author. However, a problem arises when multiple groups collaborate, making it so that there is not one “senior” author. In cases where two “senior” authors made equal contributions, I recommend indicating that with a footnote (e.g., “these two authors contributed equally”). However, even with a footnote, a decision still needs to be made about order. I recommend that, if one person would benefit more from the last author position (say, because they are pre-tenure), that person should be listed last. If the two people are at similar career stages (or if there’s another reason why the recommendation in the previous sentence doesn’t make sense), they should flip a coin (or use some other random method) and indicate in the footnote that that’s how the decision was made. If the collaboration results in more than one contribution with equal last authorship, the authors could alternate in an ABBA sequence as a means of balancing out equal contributions over time. (These same general guidelines could be applied in cases of shared first authorship as well.) Given the continued potential for confusion regarding what is conveyed by authorship order – especially in more complicated situations arising from collaborations between multiple research groups – and given the high stakes of tenure and promotion decisions, it might be advisable to include a short paragraph in the dossier that describes the authorship system that was used (e.g., a first-last author emphasis system) and noting exceptions (e.g., for a high profile paper based on work done in several different research groups).

When making decisions related to authorship, it is also important to keep in mind that individuals likely have biases that might influence who is viewed as “senior” and that this might impact views on who should be last author on a manuscript. A recent study found that only ∼25% of last authors in the journal *Functional Ecology* were women (Fox et al. 2016). It is likely that at least some of this pattern can be attributed to women being more likely to leave science, leading to fewer women as senior authors (Fox et al. 2016). At the same time, the same biases that contribute to women disproportionately leaving science (e.g., Moss-Racusin et al. 2012) might also influence decisions regarding which author is viewed as “senior” (and, therefore, in the emphasized last author position). Thus, in addition to recommending that authorship discussions begin with the default of having the senior author as last author, I also recommend that, when thinking about who is the senior author, people should be aware of potential biases (such as those related to gender or race/ethnicity) that might influence who they view as “senior”.

Of the papers published in 2016 that were examined for this study, 84% had the first author as the corresponding author. Based on the survey results, most people will assume that this person “uploaded the files, managed the revisions and wrote the response to reviewers, and took responsibility for the paper after publication”, but 19% will think it simply means that that is the person who uploaded the files. Thus, there is substantial variation in how people view corresponding authorship, including whether it is viewed as something that indicates something larger about responsibility for the work reported in the manuscript. Further work on this topic – especially studies that collect qualitative data on the topic – would be useful for understanding current views on corresponding authorship. One potential focus for such studies is whether corresponding authorship is perceived differently depending on whether the corresponding author is the first or last author, as was found in a survey of medical school department chairs (Bhandari et al. 2014). Based on the combination of poll results and current corresponding authorship practices, a reasonable starting point for discussions of authorship on ecology articles would be to have the lead author be the corresponding author on a paper noting that, in doing so, many readers will assume that means that person is taking full responsibility for the paper.

One important conclusion from this study is that there is much more work to be done on this topic. This study has several limitations – most notably relying heavily on an online survey of blog readers to understand current views on last and corresponding authorship. One problem arising from this approach is that it almost certainly skewed the age distribution of respondents (as compared to the age distribution of ecologists as a whole). In addition, the survey design (multiple choice questions) doesn’t allow insight into what factors people were weighing as they decided between different options, nor into what caveats they may have wished to add as they chose a response. Moreover, people likely varied in terms of how they interpreted some of the options (e.g., does full responsibility simply mean that is the person who handles all the requests for more information, or does it mean that, if a major problem was found with the paper, that person would take full responsibility for it?) This topic would benefit greatly from study by someone with training in social science methods, including survey design and qualitative research methods. Such work could provide further insights into the factors that influence individual author’s decisions regarding last and corresponding authorship, as well as the ways in which search committees, tenure and promotion committees, and others view authorship.

Authorship carries with it both credit and responsibility, and the order of authorship can convey information about how much credit and responsibility an author of a multi-authored paper deserves. However, because of variation across fields and over time, what is indicated by last authorship and corresponding authorship is not necessarily clear. My analyses indicate that most ecologists view the last author as the “senior” author on a paper (that is, the head of the group where the majority of the work was carried out), that the first author tends to be the corresponding author on ecology papers, and that most ecologists interpret corresponding authorship as taking full responsibility for a paper. Thus, in addition to agreeing with earlier calls to discuss authorship early and often (Weltzin et al. 2006), I suggest that those discussions have as their starting point that the last author is the senior author and the first author is the corresponding author. Collaborations between multiple groups have the potential to be trickier, but the general guidelines given above can help resolve ties that arise from equal contributions.

## Acknowledgments

This poll was confirmed as exempt from ongoing IRB review (UMich IRB #: HUM00114140). The poll was developed with input from Alex Bond, Linda Campbell, Kathy Cottingham, and Andrea Kirkwood, who all helped me think through what to ask about and how to phrase the questions and answer options. Thanks to Rayna Harris for introducing me to the Likert package and providing code for the initial version of Figure 5, to Charles Fox for letting me know that data on authorship could be downloaded from Web of Science, and to Stephen Heard and an anonymous reviewer for helpful comments on an earlier draft of the manuscript.

